# Sensorimotor expectations bias motor resonance during observation of object lifting: The causal role of pSTS

**DOI:** 10.1101/839712

**Authors:** Guy Rens, Vonne van Polanen, Alessandro Botta, Mareike A. Gann, Jean-Jacques Orban de Xivry, Marco Davare

**Author notes:** **Corresponding Author:** Guy Rens, The Brain and Mind Institute, University of Western Ontario, Ontario N6A 3K7, Canada.

## Abstract

Transcranial magnetic stimulation (TMS) studies have highlighted that corticospinal excitability (CSE) is increased during observation of object lifting, an effect termed ‘motor resonance’. This facilitation is driven by movement features indicative of object weight, such as object size or observed movement kinematics. Here, we investigated in 35 humans (23 females) how motor resonance is altered when the observer’s weight expectations, based on visual information, do not match the actual object weight as revealed by the observed movement kinematics. Our results highlight that motor resonance is not robustly driven by object weight but easily masked by a suppressive mechanism reflecting the correctness of the weight expectations. Subsequently, we investigated in 24 humans (14 females) whether this suppressive mechanism was driven by higher-order cortical areas. For this, we induced ‘virtual lesions’ to either the posterior superior temporal sulcus (pSTS) or dorsolateral prefrontal cortex (DLPFC) prior to having participants perform the task. Importantly, virtual lesion of pSTS eradicated this suppressive mechanism and restored object weight-driven motor resonance. In addition, DLPFC virtual lesion eradicated any modulation of motor resonance. This indicates that motor resonance is heavily mediated by top-down inputs from both pSTS and DLPFC. Altogether, these findings shed new light on the theorized cortical network driving motor resonance. That is, our findings highlight that motor resonance is not only driven by the putative human mirror neuron network consisting of the primary motor and premotor cortices as well as the anterior intraparietal sulcus, but also by top-down input from pSTS and DLPFC.

**Significance Statement:** Observation of object lifting activates the observer’s motor system in a weight-specific fashion: Corticospinal excitability is larger when observing lifts of heavy objects compared to light ones. Interestingly, here we demonstrate that this weight-driven modulation of corticospinal excitability is easily suppressed by the observer’s expectations about object weight and that this suppression is mediated by the posterior superior temporal sulcus. Thus, our findings show that modulation of corticospinal excitability during observed object lifting is not robust but easily altered by top-down cognitive processes. Finally, our results also indicate how cortical inputs, originating remotely from motor pathways and processing action observation, overlap with bottom-up motor resonance effects.

## Introduction

Over two decades ago, Fadiga et al. (1995) demonstrated the involvement of the human motor system in action observation: By applying single pulse transcranial magnetic stimulation (TMS) over the primary motor cortex (M1), they revealed that corticospinal excitability (CSE) was similarly modulated when observing or executing the same action. In line with the mirror neuron theory, they argued that the motor system could be involved in action understanding through a bottom-up mapping (‘mirroring’) of observed actions onto the cortical areas that are involved in their execution (for a review see: Rizzolatti et al., 2014). Consequently, action observation-driven modulation of CSE has been termed ‘motor resonance’.

Recently, TMS studies in humans substantiated that motor resonance reflects movement features within observed actions. For example, Alaerts et al. (2010a, 2010b) demonstrated that motor resonance during observation of object lifting is modulated by observed features indicative of object weight, such as intrinsic object properties (e.g. size), muscle contractions and movement kinematics. Specifically, CSE is increased when observing lifts of heavy compared to light objects. Interestingly, Alaerts et al. (2012) also demonstrated weight-driven motor resonance is already present during the observed reaching phase, suggesting an underlying predictive mechanism as well.

However, motor resonance does not seem to be robust. For instance, Buckingham et al. (2014) demonstrated, using the size-weight illusion, that CSE modulation is driven by object size when observing skilled but not erroneous lifts. In addition, Senot et al. (2011) demonstrated that object weight-driven motor resonance is eradicated when objects with identical appearance but different weights are labelled the same. Last, Tidoni et al. (2013) demonstrated that motor resonance is altered by the intentions conveyed by the observed person: CSE is increased when observing deceptive lifts compared to truthful ones. Although the above studies experimentally manipulated the information participants perceived, they could not investigate whether the participants’ expectations changed and to which extent this affected CSE modulation.

In the present study, we investigated whether the observer’s expectations alter motor resonance by manipulating the experimental context. We asked participants to perform an object lifting task in turns with an actor. One group performed the task on objects with congruent only size-weight relationship (i.e. big-heavy or small-light objects; ‘congruent objects’) whereas the other group lifted both congruent and ‘incongruent objects’ (i.e. big-light or small-heavy objects). Based on Alaerts et al. findings (2010b, 2012), we hypothesized that motor resonance would be driven (i) by the intrinsic object properties (i.e. size) before observed object lift-off and (ii) by the movement kinematics (i.e. actual object weight) after observed lift-off. However, our results revealed that, for the group lifting both congruent and incongruent objects, CSE was decreased when observing lifts of congruent objects, irrespective of the object’s size and weight. In contrast, CSE was increased when observing lifts of incongruent objects, again irrespective of size and weight. As such, motor resonance was not driven by size or weight but rather by congruence of the objects’ size-weight relationship.

We carried out a second experiment to investigate whether object weight-driven motor resonance during observed lifting was suppressed by top-down inputs to the motor system: Another group of participants performed the same task on the congruent and incongruent objects after receiving a virtual lesion of either the posterior superior temporal sulcus (pSTS) or dorsolateral prefrontal cortex (DLPFC). We opted for these areas considering their involvement in understanding intentions and motor goals [DLPFC: Miller and Cohen, (2001), Kilner (2012); pSTS: Nelissen et al. (2011)] and in recognizing action correctness [DLPFC: Pazzaglia et al. (2008); pSTS: Pelphrey et al. (2004)]. Based on evidence that pSTS is reciprocally connected with the anterior intraparietal cortex (AIP) (Nelissen et al. 2011) and DLPFC with the ventral premotor cortex (PMv) (Badre and D’Esposito 2009), which are considered key nodes for driving motor resonance (Rizzolatti et al. 2014), we hypothesized that virtual lesion of either region would release the ‘suppression’ and restore weight-driven motor resonance.

## Methods

### Participants

68 participants were recruited from the student body of KU Leuven (Belgium) and divided into four groups. 9 individuals were excluded prior to participation based on screening for TMS (Rossi et al. 2011) and/or MRI safety (checklist of local hospital: UZ Leuven). For experiment 1, 18 individuals (12 females; mean age ± SEM = 23.78 ± 0.12 years) were assigned to the control group and 17 (11 females; mean age ± SEM = 24.63 ± 0.14 years) to the baseline group. For the second experiment, 24 individuals were separated into two groups. Prior to performing the experimental task, 12 participants received virtual lesioning of DLPFC (5 females; mean age ± SEM = 24.04 ± 0.23 years) and the other 12 received virtual lesioning of pSTS (9 females; mean age ± SEM = 22.54 ± 0.18 years). The Edinburgh Handedness Questionnaire (Oldfield 1971) revealed that all participants were strongly right-handed (> 90). All participants had normal or corrected-to-normal vision, were free of neurological disorders and had no motor impairments of the right upper limb. Participants gave written informed consent and were financially compensated for their time. The protocol was in accordance with the Declaration of Helsinki and was approved by the local ethical committee of KU Leuven, Belgium (Project s60072).

### Experimental set-up

#### Experimental task

Subject and actor were comfortably seated opposite to each other in front of a table (for the experimental set-up see: figure 1A). Participants were required to grasp and lift the manipulandum (see: *‘*acquisition of force data’) that was placed in front of them in turns with the actor. As such, one trial consisted of one lifting action performed by either the actor (‘actor trial’) or the participant (‘participant trial’). Prior to the start of the task, participants received two practice trials on the objects with a congruent size-weight relationship (‘congruent objects’) but not on those with an incongruent relationship (‘incongruent objects’; for an explanation see: ‘acquisition of force data’). Participants also received the following instructions beforehand: (1) Lift the manipulandum to a height of approximately 5 cm at a smooth pace that is natural to you. (2) Only place thumb and index finger on the graspable surfaces (precision grip). (3) The cube in your trial always matches the cube in the actor’s preceding trial both in size and weight. As such, participants always lifted the exact same cube as the actor did in the preceding trial and could rely on lift observation to estimate object weight for their own trials (Rens and Davare, 2019). Finally, both participants and actor were asked to place their hand on a predetermined location on their side of the table to ensure consistent reaching throughout the experiment. Reaching distance was approximately 25 cm and required participant and actor to use their entire right upper limb to reach for the manipulandum. Lastly, participants were not informed about the incongruent objects prior to the start of the experiment.

**Figure 1.**
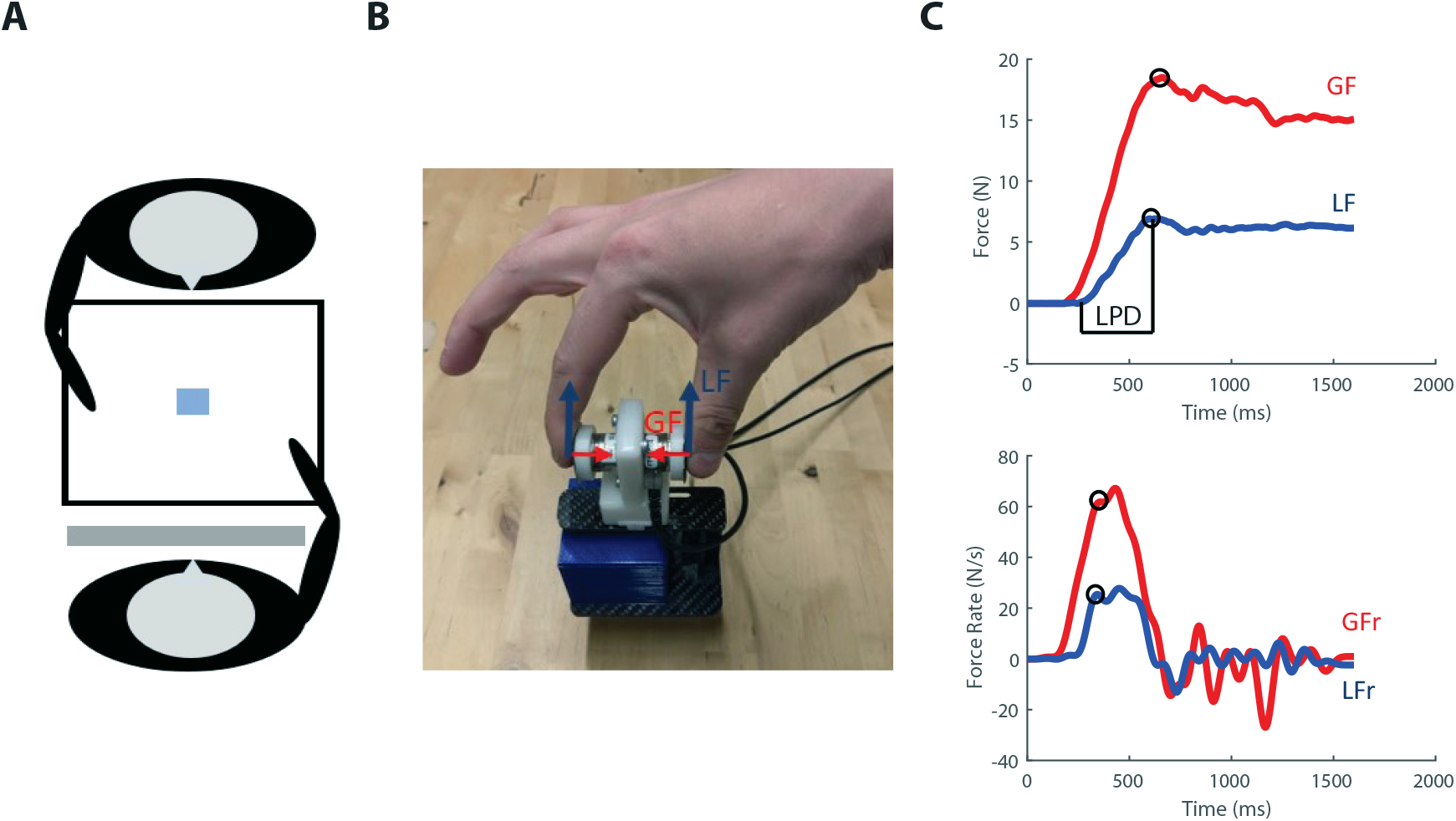
Experimental set-up. **A.** Representation of the experimental set-up: the participant and actor were seated opposite to each other in front of a table on which the manipulandum was positioned. A switchable screen was placed in front of the participant’s face. **B.** Photo of the grip-lift manipulandum used in the experiment. Load force (LF: blue) and grip force (GF: red) vectors are indicated. **C.** GF and LF typical traces (upper) and their derivatives (lower) for a skilled lift. Circles denote first peak values used as parameters. Loading phase duration (LPD) was defined as the delay between object contact (GF > 0.20 N) and object lift off (LF > 0.98*object weight).

For experiment 1 (control and baseline groups), each trial performed by the actor or the participant was initiated with a neutral sound cue (‘start cue’). For experiment 2 (DLPFC and pSTS groups), we removed the start cue as we applied TMS during participant trials as well (see the ‘TMS procedure and EMG recording’ section for the stimulation conditions; see the ‘Experimental groups’ paragraph below for the inter-group differences). Accordingly, participants in experiment 2 were instructed to consider the TMS pulse as the start cue and only initiate their movement after TMS was applied. For all groups, trials lasted 4 s to ensure that participants and actor had enough time to reach, grasp and lift the manipulandum smoothly at a natural pace. Inter-trial interval was approximately 5 s during which the cuboid in the manipulandum could be changed. A transparent switchable screen (Magic Glass), placed in front of the participant’s face, became transparent at trial onset and turned back to opaque at the end of the trial. The screen remained opaque during the inter-trial interval to ensure participants had no vision on the cube switching. The actor always performed the act of changing the cuboid before executing his trials (even if the same cube would be used twice in a row). This was done to ensure that participants could not rely on sound cues to predict cube weight in the actor’s upcoming trial. Switching actions were never performed before participant trials as they were explained that their cube would always match that of the actor.

#### Experimental procedure

All participants performed the object lifting task in a single session (‘experimental session’). Moreover, participants of experiment 2 underwent prior MRI scanning (session duration: 30 min) on a different day. At the start of the experimental session (start of scanning session for the participants of experiment 2), participants gave written informed consent and were prepared for TMS stimulations as described below. Afterwards participants performed the experimental task (for the amount of trials per group see table 1). Experimental sessions lasted 60 minutes for the control group and 90 minutes for the baseline, DLPFC and pSTS groups. Differences in session duration between the groups resulted from differences in TMS preparation and the amount of trials per group (see below).

**Table 1.**
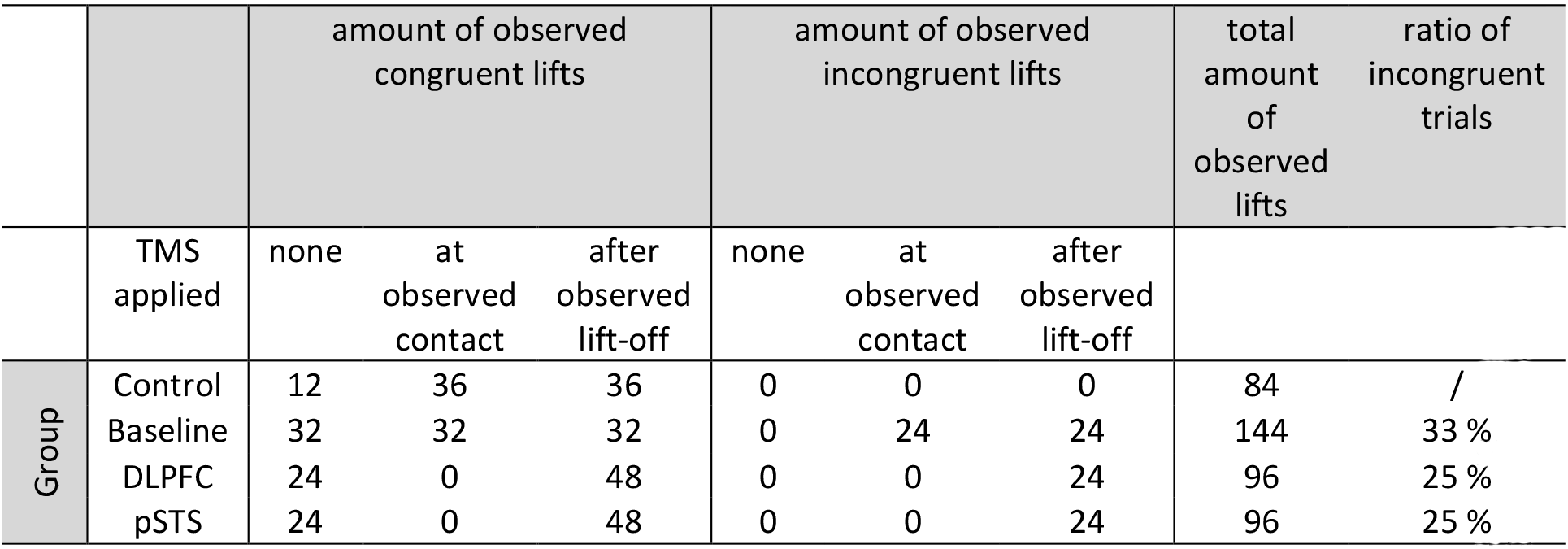
Distribution of trials per group. For each group, the amount of observed trials for each TMS condition is presented. **Amount of observed congruent lifts**: Half the amount of observed congruent lifts consisted of the big-heavy object and the other half of the small-light object. **Amount of observed incongruent lifts**: Half the amount of observed incongruent lifts consisted of the big-light object and the other half of the small-heavy object. Please note that participants lifted the objects themselves after lift observation. As such, the total amount of trials (observation and execution) is double the amount of the ‘observed trials’.

#### Experimental groups

In experiment 1, we wanted to investigate whether the presence of incongruent objects alters motor resonance. To do so, we divided participants into two groups: the control and baseline group. Participants in the control group were only exposed to the congruent objects. In contrast, participants in the baseline group lifted both the congruent and incongruent objects during the task.

In experiment 2, we wanted to investigate how pSTS and DLPFC are causally involved in mediating the suppressive mechanism revealed in experiment 1. Participants performed the same task as the baseline group of experiment 1 (that is, interacting with both the congruent and incongruent objects) after receiving a virtual lesion over either pSTS or DLPFC.

#### Trial amount per experimental group

First, we initially considered the incongruent objects to be pivotal for investigating how motor resonance is driven by expected and actual object weight. We decided on 12 trials per object condition for the incongruent objects (12 for small-heavy and 12 for big-light; 24 in total for both incongruent objects combined) based on Senot et al. (2011). Their study, is to our knowledge, one of the few that investigated motor resonance during observation of ‘live’ (no video recordings) observation of object lifting. As they found consistent results using 10 trials per condition, we decided to include two more due to our experimental task. These two extra trials were intended to serve as a buffer for potential errors made by the actor or the participants.

Second, we decided to use unequal proportions of congruent and incongruent objects based on Alaerts et al. (2012). They demonstrated that, during lift observation, the observer’s motor system predictively encodes object weight during the observed reaching phase. However, it is important to note that they used a blocked design, enabling participants to anticipate object weight even though the objects were visually identical. Considering that we did not want to rely on a blocked design but a pseudo-randomized one, we argued that unequal proportions would cause participants to expect that size was indicative of weight, causing motor resonance to be driven by these size-driven weight expectations at observed contact. In contrast, we argued that, if presented with equal proportions, participants would entirely ignore the size cue (as it could indicate either weight) eradicating motor resonance at observed object contact.

Third, we initially wanted 25 % of trials to be incongruent for all groups interacting with both congruent and incongruent objects. However, this was not feasible for the baseline group as this would cause their behavioral task to last twice as long compared to the other groups. Accordingly, we increased the amount of incongruent trials to 33 % for the baseline group. This proportion was selected based on Pavone et al. (2016). They showed that neural activity, recorded with EEG, is different when observing correctly (70% of trials) and incorrectly (30% of trials) executed grasping actions in virtual reality. Importantly, this proportional difference between our baseline group (33 %) on one side and the DLPFC and pSTS groups (25 %) on the other side should not have affected motor resonance differently: Pezzetta et al. (2018) demonstrated, using EEG, that observed errors rather than their probability elicit typical error-related cortical activation. Last, for all groups the amount of congruent trials was defined with the intent of maintaining these proportions of incongruent trials.

Fourth, as our findings for the baseline group showed that motor resonance was not modulated at observed contact, we decided to remove this TMS timing condition for the DLPFC and pSTS groups. This was done to ensure that the behavioral task was completed before the disruptive effects of cTBS, lasting approximately one hour (Huang et al., 2005), ran out.

Fifth, to investigate whether TMS during lift observation did not interfere with the participants’ lift planning, we included a non-TMS condition for the congruent objects (33 % of congruent trials amount).

Last, we included the control experiment due to our unanticipated findings in the baseline group. As our baseline group findings showed that TMS did not interfere with predictive lift planning, we decided to reduce the amount of non-TMS trials. We decided to include two more trials (18 in total) for each (congruent) condition compared to the baseline experiment. These trials were intended to serve as a buffer for potential errors made by the actor or the participant and to ensure we minimally had 16 correct congruent trials.

#### Object lifting sequences

A unique pseudo-randomized object lifting sequence was generated for each participant of each group using a custom-written MATLAB script. For the baseline group, this sequence was divided over four experimental blocks. For participants in the control, DLPFC and pSTS group, this sequence was divided over two experimental blocks. Participants received a short break between experimental blocks. Pseudo-randomization was based upon the following criteria: (i) Within each experimental block, objects of the same condition were presented an equal amount of times (e.g. In a given experimental block, half the amount of congruent objects were big-heavy whereas the other half was small-light). (ii) Each object for each TMS timing was presented an equal amount of times in each experimental block (e.g. For the baseline group, lift observation of the big-heavy object when TMS was applied at observed object contact was presented four times in each of the four experimental blocks). (iii) Each experimental block of the baseline, DLPFC and pSTS groups could not start with an incongruent trial. (iv) Two incongruent trials were separated by at least one congruent trial. (v) Half the amount of incongruent trials, in a given experimental block, were performed in the first half of that experimental block and the other half of incongruent trials in the second half of that experimental block.

### Acquisition of force data

A grip-lift manipulandum consisting of two 3D force-torque sensors was attached to a custom-made carbon fiber basket in which different objects could be placed (for an image of the manipulandum see: figure 1B). The total weight of the manipulandum was 1.2 N. The graspable surface (17 mm diameter and 45 mm apart) of the force sensors was covered with fine sandpaper (P600) to increase friction. For the present experiment, we used four 3D-printed objects. The large objects (cuboids) were 5×5×10 cm in size whereas the two small ones (cubes) measured 5×5×5 cm. Two of the objects, one small and one large, were filled with lead particles so each of them weighted 0.3 N. The other two were filled with lead particles until each of them weighted 5 N. Combined with the weight of the manipulandum, the light and heavy objects weighted 1.5 and 6.3 N respectively. Importantly, using these four objects, we had a two by two design with size (small or big) and weight (light or heavy) as factors. In addition, this design allowed us to have two objects that were ‘congruent’ in size and weight (large objects are expected to be heavier than smaller ones of the same material) and two ‘incongruent’ objects for which this size-weight relationship was inversed (Baugh et al. 2012). To exclude any visual cues indicating potential differences between the same-sized objects, they were hidden under the same paper covers. In the present study, we used two ATI Nano17 F/T sensors (ATI Industrial Automation, USA). Both F/T sensors were connected to the same NI-USB 6221 OEM board (National Instruments, USA) which was connected to a personal computer. Force data was acquired at 1000 Hz using a custom-written Labview script (National Instruments, USA). Lastly, one of the authors G. Rens served as the actor in both experiment 1 and 2.

### TMS procedure and EMG recording

#### General procedure

For all groups, electromyography (EMG) recordings were performed using Ag-AgCl electrodes which were placed in a typical belly-tendon montage over the right first dorsal interosseous muscle (FDI). A ground electrode was placed over the processus styloideus ulnae. Electrodes were connected to a NL824 AC pre-amplifier (Digitimer, USA) and a NL820A isolation amplifier (Digitimer, USA) which in turn was connected to a micro140-3 CED (Cambridge Electronic Design Limited, England). EMG recordings were amplified with a gain of 1000 Hz, high-pass filtered with a frequency of 3 Hz, sampled at 3000 Hz using Signal software (Cambridge Electronic Design Limited, England) and stored for offline analysis. For TMS stimulation, we used a DuoMAG 70BF coil connected to a DuoMAG XT-100 system (DEYMED Diagnostic, Czech Republic). For M1 stimulation, the coil was tangentially placed over the optimal position of the head (hotspot) to induce a posterior-anterior current flow and to elicit motor evoked potentials (MEPs) in right FDI. The hotspot was marked on the scalp of each participant. Stimulation intensity (1 mV threshold) for each participant was defined as the lowest stimulation intensity that produced MEPs greater than 1 mV in at least four out of eight consecutive trials when stimulating at the predetermined hotspot. Last, the control group and baseline group received 12 stimulations at the 1 mV threshold before and after the experiment to have a baseline measure of resting CSE. Moreover, for the baseline group, we also recorded a baseline measure of resting CSE halfway through the experiment (i.e. when participants had performed half of the experimental blocks) as their experimental session lasted 30 min longer.

#### Stimulation during the experimental task

For the control and baseline group, single-pulse TMS over M1, for probing CSE, was applied during the actor trials at two different timings: at observed object contact and 300 ms after observed object lift-off (see ‘Data processing’ for definitions of object contact and lift-off). Participants did not receive stimulations during their trials (i.e. participant trials).

For the DLPFC and pSTS groups, single-pulse TMS, over M1 for probing CSE, was applied during both the actor and participant trials. During observation we only applied single-pulse TMS during the observed lifting phase, and not at observed contact for two reasons: (1) The results from experiment 1 indicated that CSE was primarily modulated after observed object lift-off and (2) because of the time constraints related to the duration of the after-effects caused by cTBS (Huang et al. 2005), which are limited to about an hour. During participant trials, single-pulse TMS was applied 400 ± 100 ms (jitter) after object presentation. As participants were instructed to only start lifting after receiving the stimulation, it was applied during movement planning and not execution. We did not stimulate the control and baseline groups during lift planning because, initially, we were only interested in motor resonance. We then included these stimulations in experiment 2, because we wanted to investigate the effect of a virtual lesion of DLPFC or pSTS on CSE modulation during motor planning and whether these effects would be different from those during action observation. Finally, in experiment 1 (control and baseline groups) we did not use neuro-navigation but relied on the hotspot mark on the scalp to apply single-pulse TMS over M1 during the experiment. In contrast, for experiment 2 (DPLFC and pSTS groups) we used neuro-navigation for applying cTBS over these regions but also for maintaining the same coil positioning and orientation when applying single-pulse TMS over M1 during the experiment. Accordingly, for experiment 2, the hotspot was determined using the same procedures as in experiment 1, although the single-pulse TMS stimulations over M1 during the experiment were neuro-navigated. However, this should not have affected the validity of our between-group differences (for example see: Jung et al., 2010).

#### Additional procedures for experiment 2

After defining the 1 mV threshold, we defined the active motor threshold (aMT) as the lowest stimulation intensity that produced MEPs that were clearly distinguishable from background EMG during a voluntary contraction of about 20 % of their maximum using visual feedback. Before the experimental task, participants received cTBS over either DLPFC or pSTS. cTBS consisted of bursts of 3 pulses at 50 Hz, repeated with a frequency of 5 Hz and at an intensity of 80 % of the aMT for 40 s (600 pulses in total). It has been considered that this type of repetitive stimulation disrupts activity within the stimulation region for a period up to 60 minutes (Huang et al. 2005). Consequently, it has often been termed a ‘virtual lesion’. In experiment 2, we also collected resting CSE before cTBS. As such, we recorded three resting CSE measurements, i.e. pre-cTBS, pre-task (5 minutes after cTBS ended and just before the start of the experimental task) and post-task. To ensure that cTBS was applied on the desired stimulation area, a high-resolution structural T1-weighted anatomical image of each participant was acquired with a magnetization-prepared rapid-acquisition gradient-echo (MPRAGE) sequence (Philips Ingenia 3.0T CX, repetition time/echo time = 9.72/4.60 ms; voxel size = 1.00 × 1.00 × 1.00 mm^3^; field of view = 256 × 256 × 192 mm^3^; 192 coronal slices) which was co-registered during the experiment with the fiducial landmarks using a Brainsight TMS neuronavigation system (Rogue Research, Canada).

DLPFC was anatomically identified following Mylius et al. (2013). Briefly, we identified the superior and inferior frontal sulci as the superior and inferior borders of the middle frontal gyrus (MFG). The posterior border was defined as the precentral sulcus and the frontal one as the anterior termination of the olfactory sulcus in the coronal plane. Lastly, the MFG was divided equally into three parts and the separating line between the anterior and middle thirds was defined as the DLPFC (for full details see: Mylius et al., 2013). We always defined DLPFC within the middle frontal sulcus (MFS). This allowed us to consistently target the MFS using the same coil orientation across participants. Coil orientation was perpendicular to the MFS with the handle pointing downwards. pSTS was anatomically defined following Cattaneo et al. (2010) and Arfeller et al. (2013) as the middle between the caudal and rostral ends of the ascending branch of STS, just below the intraparietal sulcus. Coil orientation was perpendicular to pSTS with the handle pointing downwards. The means ± SEM of Talaraich coordinates for these sites were as follows: left DLPFC: X = −38.14 ± 0.93, Y = 23.53 ± 1.64, Z = 32.29 ± 0.80; left pSTS: X = −54.03 ± 1.09, Y = −49.86 ± 1.32, Z = 9.35 ± 1.22 as estimated on the cortical surface (For stimulation locations see: figure 2) which are in line with previous studies [left DLPFC: X = −42.17 ± 5.07, Y = −33.73 ± 5.73, Z = 32.36 ± 6.17 Mylius et al. (2013); left pSTS: X = −51.6 ± 3.6, Y = −43.2 ± 7.1, Z = 7.1 ± 6.4 Arfeller et al. (2013)].

**Figure 2.**
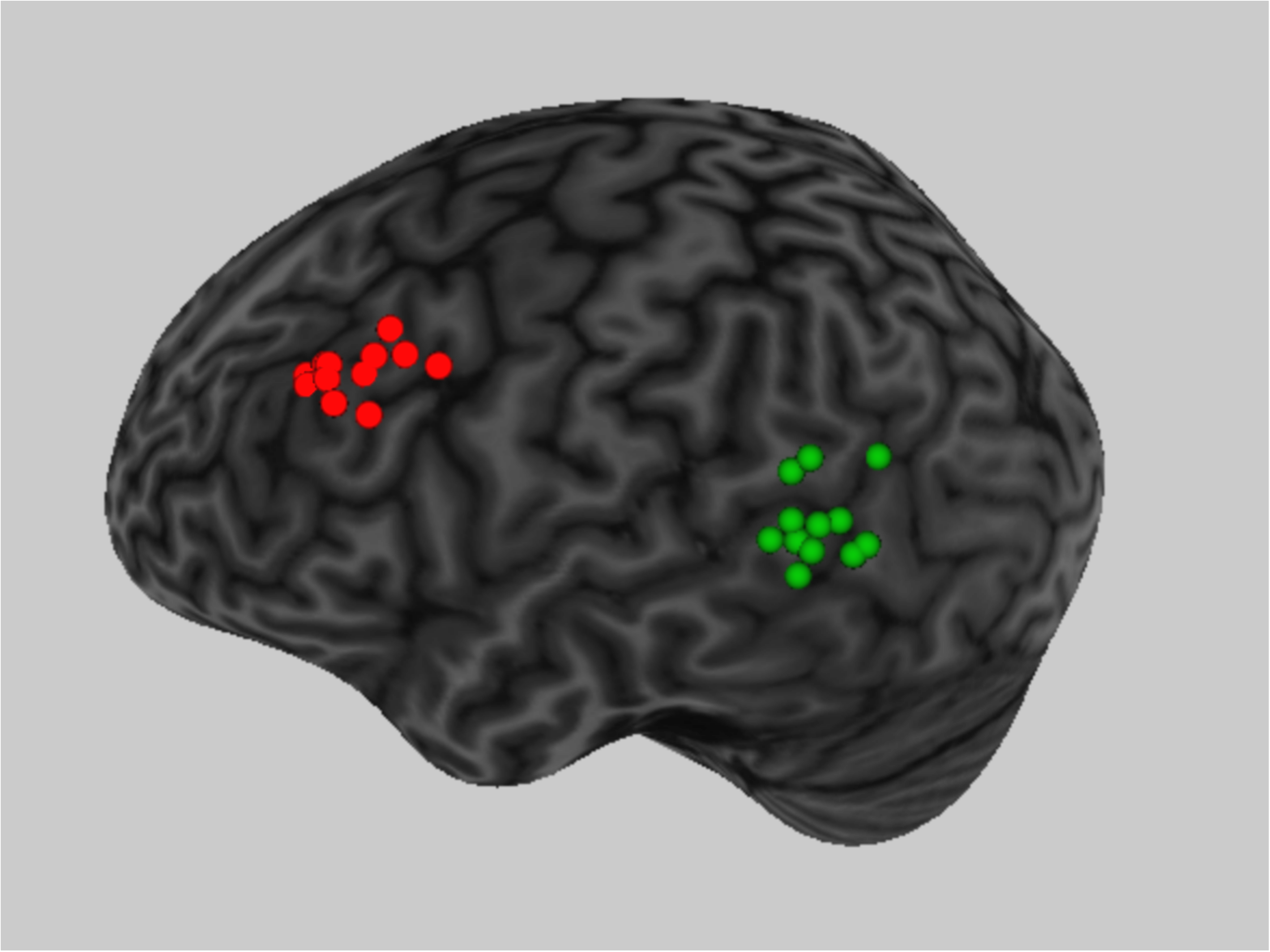
Stimulation sites. Anatomical locations where cTBS was applied for each individual subject of the DLPFC (red) and pSTS (green) groups.

### Data processing

#### Force data

Data collected with the F/T sensors were low-pass filtered with a fifth-order Butterworth filter (forces cut-off frequency: 30 Hz, force rates cut-off frequency: 15 Hz). A custom script was written in MATLAB to compute the following variables: (1) Grip (GF) and load (LF) forces, which were defined as the exerted force perpendicular and tangential to the normal force, respectively (figure 1B). GF and LF were computed as the sum of the respective force components exerted on both sensors. Additionally, grip and load force rates (GFr and LFr) were computed by taking the first derivative of GF and LF respectively. We report not GF and LF but their rates (figure 1C) as it has been demonstrated that force rate parameters are a reliable indicator of predictive force scaling (Gordon et al. 1991; R. S. Johansson and Westling 1988). For analyses purposes of the force parameters, we decided to use the first peak grip and load force rate values after object contact that were at least 30 % of the maximum peak rate. This threshold was used to exclude small peaks in the force rates due to noise or small bumps caused by lightly contacting the F/T sensors. In addition, we decided to use the first peak force rate values as later peak values might be contaminated with feedback mechanisms and not reflect predictive force planning (Castiello 2005; Rens and Davare 2019). Accordingly, using the peak force rates enabled us to investigate whether participants scaled their fingertips forces differently for the incongruent and congruent objects. Besides peak force rates, we also report the loading phase duration (LPD) which was defined as the latency between object contact and lift off. Object contact and lift-off were defined as the time points when GF exceeded 0.2 N and LF exceed 0.98 × object weight (figure 1C), respectively (please note that these definitions were used for timing the TMS stimulation during lift observation; see: ‘TMS procedure and EMG recording’). In addition, GF and LF were required to stay above these thresholds for at least 200 ms. We included LPD as it is considered an estimator of the lifting speed (e.g. the shorter the LPD the faster the object will be lifted: Johansson and Westling, 1988a), which is a movement parameter used by participants to estimate object weight (Hamilton et al., 2007). Moreover, we could also use this parameter to investigate the participants’ lifting performance. Last, both force rate parameters and LPD were z-score normalized. For the participants, z-score normalization was done for each participant separately. For the actor, z-score normalization was also done for each ‘participant’ separately. That is, the actor’s lifting performance in one session (as observed by one participant) was z-score normalized against the data of only that session. We decided to normalize our data based on the assumption that the actor’s lifting speed might vary and this might affect the participants’ lifting speed as well. Accordingly, z-score normalization would enable us compare between-group differences.

#### EMG data

From the EMG recordings, we extracted the peak-to-peak amplitudes of the MEP using a custom-written MATLAB script. All EMG recordings were visually inspected for background noise related to muscle contractions. Moreover, trials were excluded when the MEP was visibly contaminated (i.e. spikes in background EMG) or when an automated analysis found that the average background EMG was larger than 50 μV (root-mean-square error) in a time window of 200 ms prior to the TMS stimulation. We also assessed pre-stimulation (background) EMG by calculating the root-mean-square error scores across a 100ms interval ending 50ms prior to TMS stimulation. Last, for each participant separately we excluded outliers which were defined as values exceeding the mean ± 3 SD’s. For each participant, all MEPs collected during the experimental task (but not resting measurements) were normalized with z-scores using their grand mean and standard deviation. For experiment 2, z-scoring was done for lift observation and planning separately.

### Statistical analysis

#### Corticospinal excitability during rest

To investigate within-group differences in baseline CSE, we performed repeated measures analyses of variance (ANOVA_RM_) for the control and the baseline group separately with one within-factor RESTING STATE (control: pre- and post-task; baseline; pre-task, between experimental blocks, post-task). For experiment 2, we performed a mixed ANOVA with between-factor GROUP (DLPFC or pSTS) and within factor RESTING (pre-cTBS, pre-task, post-task).

#### Within-group differences for corticospinal excitability during the experimental task

First, to investigate whether our experimental task can elicit weight-driven motor resonance effects during lift observation, we performed a ANOVA_RM_ on the control group only with within-factors CUBE (big heavy or small light) and TIMING (observed contact or after observed lift-off). To investigate whether the presence of the incongruent objects altered motor resonance, we used a general linear model (GLM; due to different effect sizes) to probe potential differences between the control and baseline groups on the congruent objects only. We used the between-factor GROUP (control or baseline) and within-factors CUBE and TIMING. Due to our findings, we followed up on this GLM with a ANOVA_RM_, only performed on the baseline group with within-factors TIMING, SIZE (big or small) and WEIGHT (heavy or light).

After these analyses on the groups of the first experiment, we investigated the potential effects of the virtual lesions of DLPFC and pSTS. For this, we performed a GLM with between-factor GROUP (baseline, DLPFC or pSTS) and within-factors SIZE and WEIGHT. As we did not stimulate the DLPFC and pSTS groups at observed contact, we could not include the within-factor TIMING. As we wanted to further explore potential within-group effects, we followed up on the GLM with separate ANOVA_RM_s for the DLPFC and pSTS groups with within-factors SIZE and WEIGHT. Finally, to explore potential differences between lift observation and planning for the groups of experiment 2, we performed a final GLM with between-factor GROUP (DLPFC or pSTS) and within-factors ACTION (observation or planning), SIZE and WEIGHT.

#### Within-group differences in background EMG during the experimental task

To ensure that differences in CSE during the behavioral task were not driven by between-condition variations in background EMG, we performed the analyses described in the preceding paragraph on the background EMG as well.

#### Force parameters of the participants

For each parameter of interest (peak GFr, peak LFr and LPD), we performed a GLM on the congruent objects only with between-factor GROUP (control, baseline, DLPFC or pSTS) and within-factor CUBE (big heavy or small light). We performed an additional GLM on the congruent and incongruent objects combined with between-factor GROUP (baseline, DLPFC or pSTS; control not included due to not using the incongruent objects) and within-factors SIZE and WEIGHT. Importantly, within-factors related to the timing of the TMS stimulation are not included here as our preliminary analyses indicated that it did not affect predictive force planning in the participants, i.e. we did not find significance for any of the relevant pairwise comparisons. Based on these findings, we decided to pool the data for TIMING and present the data as such for clarity.

#### Force parameters of the actor

For each parameter (peak GFr, peak LFr and LPD) we performed the same analyses as described in ‘Force parameters of the participants’. We did not include the within-factors related to timing as the actor was blinded to the timings during the experiment.

Last, for the GLMs we used type III sum of squares, comparisons of interest exhibiting statistically significant differences *(p ≤ 0.05)* were further analyzed using the Holm-Bonferroni test. All data presented in the text are given as mean ± standard error of the mean. All analyses were performed in STATISTICA (Dell, USA).

## Results

In the present study, we investigated how motor resonance is modulated during lift observation. For this, participants performed an object lifting task in turns with an actor. The control group only lifted objects with a congruent size-weight relationship (i.e. ‘big heavy’ and ‘small light’ objects). The baseline group lifted objects with both congruent and incongruent size-weight relationships (i.e. additional ‘big light’ and ‘small heavy’ objects). The subject groups participating in experiment 2 (DLPFC and pSTS groups) used the same objects as the baseline group. Importantly, they performed the experimental task after receiving a TMS induced virtual lesion over either DLPFC or pSTS. Only relevant main and interaction effects are reported below.

### Stimulation intensities

To examine differences between stimulation intensities of the different groups, we ran two GLMs to investigate group differences in 1 mV thresholds (all groups) and aMT (DLPFC and pSTS groups only). All values are expressed as a percentage of the maximal stimulator output. As expected, the GLM failed to reveal any significant difference between groups for the 1 mV stimulation intensity (control = 61 % ± 2.62; baseline = 55.64 % ± 3.26; DLPFC = 57.54 % ± 3.26; pSTS = 50.46 % ± 3.00) *(F*_(3,48)_ = 2.39 *p = 0.08, η*^*2*^_*p*_ *= 0.13)* as well as for the aMT (DLPFC = 42.82 % ± 2.26; pSTS = 38.46 % ± 2.08) *(F*_(1,22)_ = 2.01 *p = 0.17, η*^*2*^_*p*_ *= 0.08).* Note that the degrees of freedom of the error are lower due to missing values.

We informally asked participants in experiment 2 how they perceived cTBS. In the DLPFC group, 2 out of 12 participants described cTBS as ‘uncomfortable’ whereas the other ten did not report negative sensations. In the pSTS group, five participants reported negative sensations: four reported the sensations as ‘uncomfortable’ and one as ‘painful’. Lastly, no one reported other physical adverse effects (such as dizziness or headaches) that could potentially have been related to the single pulse or cTBS stimulations.

### Corticospinal excitability at rest

#### Experiment 1

For the control (pre-task = 0.89 mV ± 0.08; post-task = 1.16 mV ± 0.22) and baseline groups (pre-block 1 = 0.61 mV ± 0.06; between-blocks = 0.79 mV ± 0.18; post-block 2 = 0.87 mV ± 0.17), both analyses provide no evidence that resting CSE changed significantly over time (non-significance of TIMING; *both F < 167, both p > 0.21, both η*^*2*^_*p*_ *< 0.09).*

#### Experiment 2

Both the main effects of GROUP, TIMING as well as their interaction effect were not significant *(all p > 0.16)* providing no evidence that resting CSE differed between groups or changed over time (DLPFC: pre-cTBS = 1.16 mV ± 0.26, pre-task = 1.53 mV ± 0.22, post-task = 1.60 mV ± 0.44; pSTS: pre-CTBS = 2.04 mV ± 0.26, pre-task = 1.60 mV ± 0.22, post-task = 2.20 mV ± 0.44).

### Background EMG during the experiment

To ensure that between-group and between-condition differences were not driven by differences in hand relaxation during lift observation and planning, we investigated potential differences in background EMG. For this we used the same statistics as described in ‘Statistical analyses - *Within-group differences for* c*orticospinal excitability during the experimental task’.* Briefly, all main and interaction effect across all analyses, except for one, were not significant *(all F < 1.99, all p > 0.18, all η*^*2*^_*p*_ *< 0.11).* The interaction effect ACTION (observe or plan lift) × SIZE (small or big) × GROUP (DLPFC or pSTS) was significant *(F = 5.14, p = 0.03, η*^*2*^_*p*_ *= 0.19).* However, the post-hoc analysis failed to reveal significant differences between any of the conditions. These findings provide no evidence that background EMG different significantly between-and within groups.

### Corticospinal excitability during the experimental task

With the control group, we investigated whether our task can elicit weight driven modulation of CSE during observed object lifting. As shown in Figure 3, the analysis substantiated the validity of our set-up: When the control group observed lifts of the big heavy object (big heavy = 0.07 ± 0.03) CSE was significantly facilitated compared to when they observed lifts of the small light object (small light = −0.08 ± 0.03; *p = 0.02*) *(main effect of CUBE: F*_*(1,17)*_ *= 6.87, p = 0.02, η*^*2*^_*p*_ *= 0.29).*

**Figure 3.**
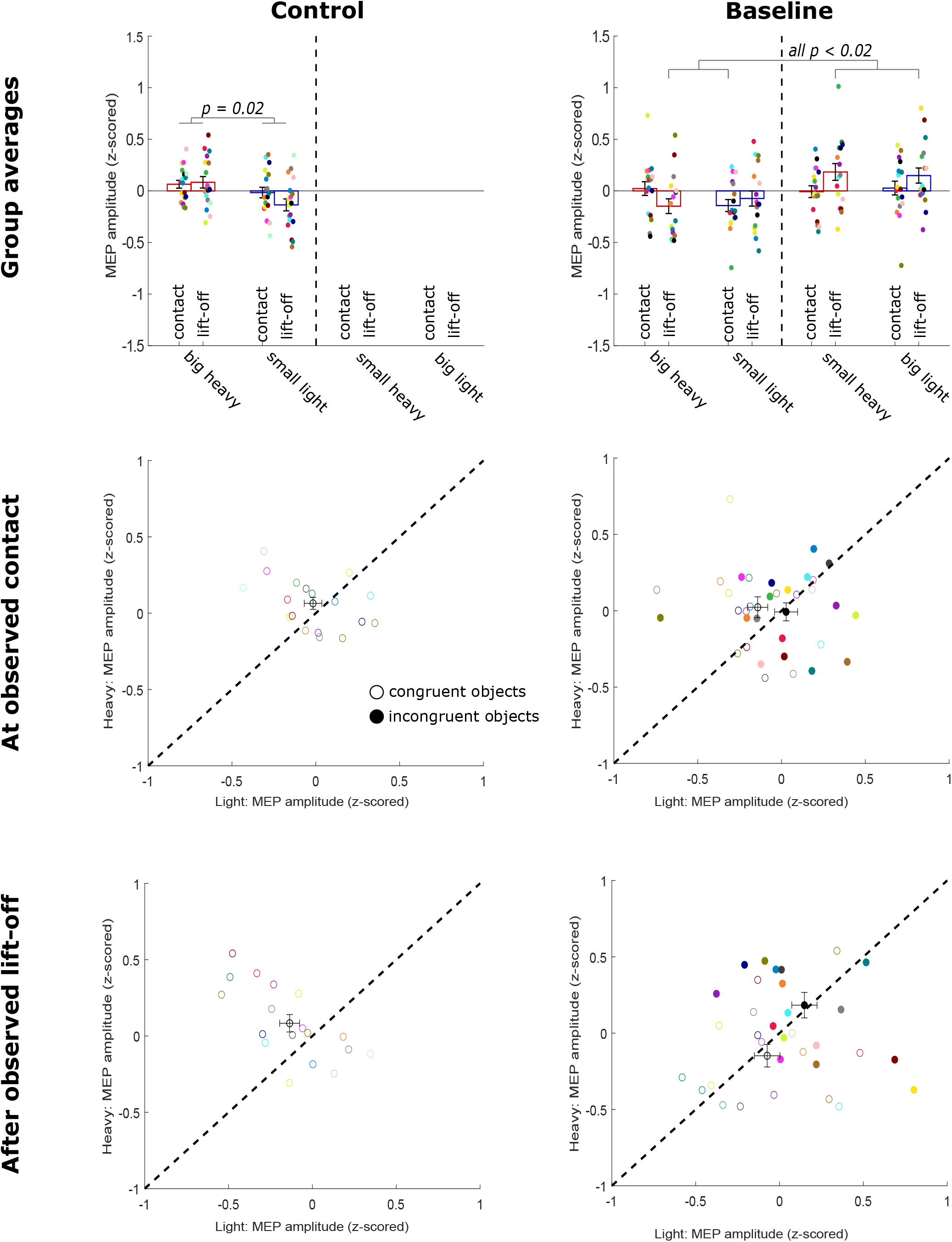
Modulation of corticospinal excitability during lift observation in the control and baseline group. **Top row.** Average MEP values (z-score) during lift observation pooled across participants for the control (left) and baseline group (right). Left and right of the dashed line on each figure represent the congruent (big heavy and small light) and incongruent objects (small heavy and big light) respectively. Red and blue indicate heavy and light weights respectively. For the control and baseline groups we used two TMS timings during observation, i.e. at observed object contact and after observed lift-off. As such, of two adjacent bars, the first and second one represent MEP values at observed contact and during observed lifting respectively. **Middle and bottom row.** 2D visualization of the average MEP values (z-scored) at observed object contact (middle) and after observed lift-off (bottom). MEP values for the heavy and light objects are shown on the y- and x-axis respectively. Each participant within each group is represented by two same colored circles (scatter). ‘Empty circles’ represent the congruent objects (MEP values for big-heavy on the y-axis and for small-light on the x-axis), ‘filled circles’ represent the incongruent objects (MEP values for small-heavy on the y-axis and for big-light on the x-axis). Black circles represent the group average ± SEM for the respective conditions. The black dashed line represents the equation y = x and indicates the line of ‘no CSE modulation’. Accordingly, scatter circles above or below the dashed line indicate that CSE, when observing lifts of heavier objects, was increased or decreased respectively. No intra-group significant differences are shown on the middle and bottom row.

Afterwards, we explored whether the presence of the incongruent objects affected motor resonance. For this, we compared the control and baseline groups for only the congruent objects. In line with our findings for the control group, CSE was significantly facilitated when observing lifts of the big heavy cube (big heavy = 0.006 ± 0.02) compared to the small light one (small light = −0.09 ± 0.03; *p = 0.04) (main effect of CUBE: F*_*(1,33)*_ *= 4.34, p = 0.04, η*^*2*^_*p*_ *= 0.12).* However, the main effect of GROUP *(F*_*(1,33)*_ *= 7.30, p = 0.01, η*^*2*^_*p*_ *= 0.18)* was significant as well: When observing lifts (of the congruent objects) CSE of the baseline group (congruent objects = −0.09 ± 0.02) was significantly more inhibited than that of the control group (congruent objects = 0.00 ± 0.02). Considering that the group averages for CSE (MEP-amplitude) are calculated using z-score normalization, these findings indicate that the presence of the incongruent objects in the baseline experiment should have inhibited CSE modulation for the congruent objects (due to negative z-score). In addition, the interaction effect CUBE × TIMING × GROUP *(F*_*(1,33)*_ *= 3.71, p = 0.06, η*^*2*^_*p*_ *= 0.10)* was borderline significant. Due to this borderline significance, we decided to explore how the presence of the incongruent objects in the baseline group affected modulation of motor resonance.

To further probe potential differences between the congruent and incongruent objects for the baseline group, we performed a separate ANOVA_RM_ on the baseline group with within-factors TIMING, SIZE and WEIGHT. Interestingly, this analysis revealed that CSE modulation in the baseline group was not driven by SIZE or WEIGHT but by ‘congruency’. As shown in Figure 3, CSE was significantly more facilitated for the small heavy object during observed lifting (mean = 0.18 ± 0.08) compared to the big heavy one during observed lifting (mean = −0.15 ± 0.07; *p = 0.01*) and the small light one at observed contact (mean = −0.14 ± 0.06; *p = 0.02*) *(interaction effect of WEIGHT × SIZE × TIMING: F*_*(1,16)*_ *= 7.54, p = 0.01, η*^*2*^_*p*_ *= 0.32).* Conversely, CSE was significantly more facilitated during observed lifting of the big light object (mean = 0.15 ± 0.08), compared to the big heavy one during observed lifting *(p = 0.03)*, and the small light one at observed contact *(p = 0.04)* (*SIZE × WEIGHT × TIMING)*. Importantly, these findings contradict our initial hypothesis: We expected that motor resonance would be driven by SIZE at observed contact and afterwards by WEIGHT during observed lifting. However, our results demonstrated that motor resonance effects driven by size or weight were ‘masked’ by a mechanism that is monitoring object congruency, i.e. monitoring a potential mismatch between anticipated and actual object weight.

With the pSTS and DLPFC groups, we investigated the potential effects of the virtual lesions on CSE modulation during lift observation. As described in ‘Statistical analysis’, we performed a GLM with between-factor GROUP (baseline, DLPFC and pSTS groups) and within-factors SIZE and WEIGHT. As shown in Figure 4, this analysis revealed that for the pSTS group, CSE was significantly facilitated when observing lifts of heavy objects, irrespective of their size (heavy objects = 0.11 ± 0.05) compared to lifts of the light ones (light objects = −0.12 ± 0.04; *p = 0.03) (interaction effect of GROUP × WEIGHT: F*_*(2,38)*_ *= 4.97, p = 0.01, η*^*2*^_*p*_ *= 0.17)*. However, this weight-driven modulation of CSE during lift observation was absent for the baseline group (due to the congruency effect as described above; heavy objects = 0.02 ± 0.04; light objects = 0.04 ± 0.03; *p = 1.00)* but was also absent for the DLPFC group (heavy objects = −0.02 ± 0.05; light objects = 0.02 ± 0.04; *p = 1.00) (GROUP × WEIGHT)*. As such, these findings indicate that weight-driven modulation of CSE during lift observation was restored for the pSTS group. However, these results do not provide any evidence that CSE was modulated after virtually lesioning DLPFC.

**Figure 4.**
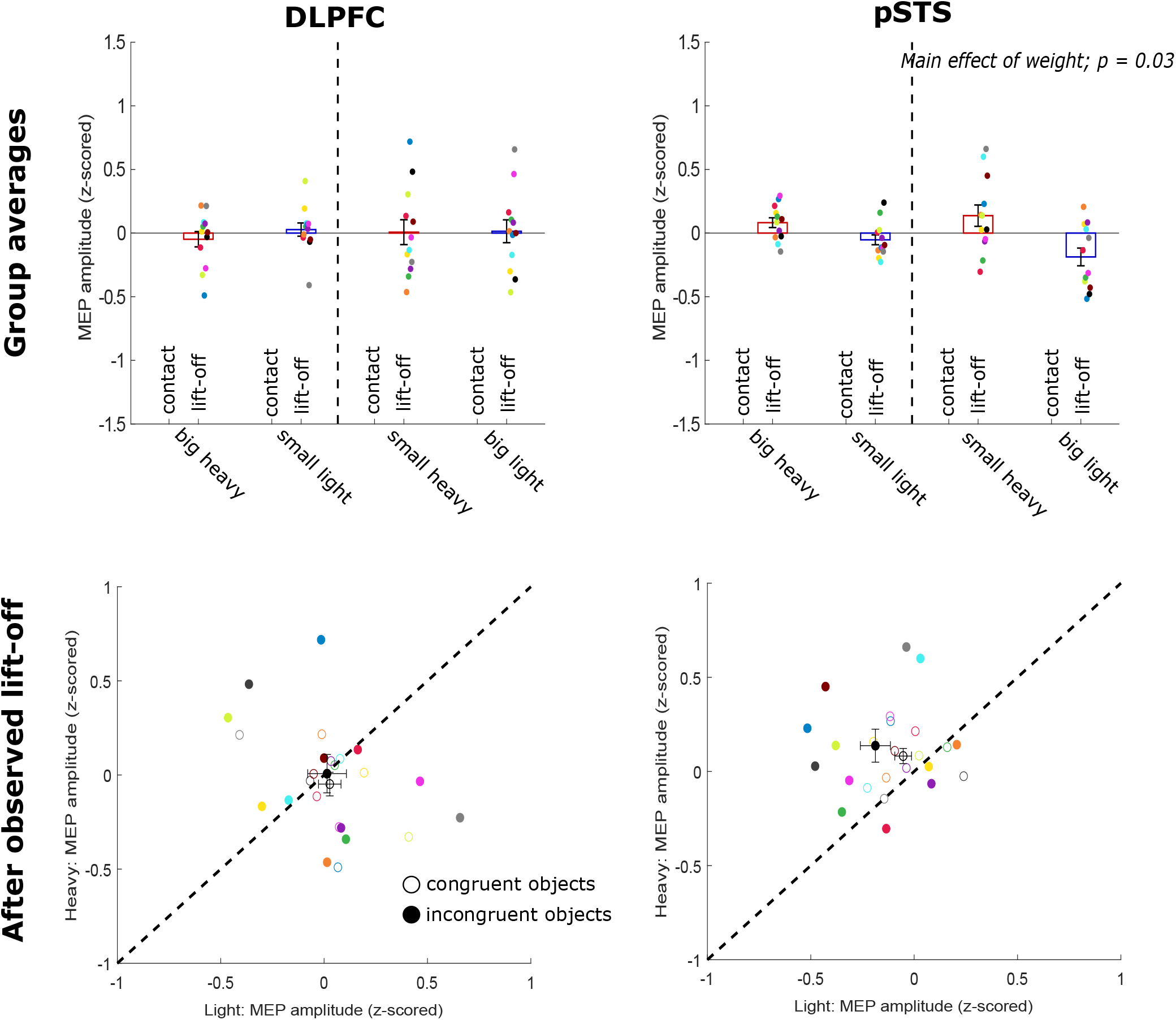
Modulation of corticospinal excitability during lift observation in the DLPFC and pSTS groups. **Top row.** Average MEP values (z-score) during lift observation pooled across participants for the DLPFC (left) and pSTS group (right). Left and right of the dashed line on each figure represent the congruent (big heavy and small light) and incongruent objects (small heavy and big light) respectively. Red and blue indicate heavy and light weights respectively. **Bottom row.** 2D visualization of the average MEP values (z-scored) after observed lift-off. MEP values for the heavy and light objects are shown on the y- and x-axis respectively. Each participant within each group is represented by two same colored circles (scatter). ‘Empty circles’ represent the congruent objects (MEP values for big-heavy on the y-axis and for small-light on the x-axis), ‘filled circles’ represent the incongruent objects (MEP values for small-heavy on the y-axis and for big-light on the x-axis). Black circles represent the group average ± SEM for the respective conditions. The black dashed line represents the equation y = x and indicates the line of ‘no CSE modulation’. Accordingly, scatter circles above or below the dashed line indicate that CSE, when observing lifts of heavier objects, was increased or decreased respectively. No intra-group significant differences are shown on the middle and bottom row.

To further investigate the WEIGHT effect in the pSTS group, we performed an additional GLM for the control and pSTS groups combined. Indeed, if weight-driven modulation of CSE during lift observation was restored by virtual lesioning of pSTS, then the pSTS group should have not differed significantly from the control group with respect to the congruent objects. For this analysis, we used the between-factor GROUP (control and pSTS) and within-factor CUBE (big heavy and small light) for TIMING being only after observed lift-off (as we did not apply TMS at observed contact in the pSTS group). Importantly, the main effect of CUBE was significant *(F*_*(1,28)*_ *= 6.43, p = 0.02, η*^*2*^_*p*_ *= 0.19)*. In line with our control group findings, CSE was significantly facilitated when observing lifts of the big heavy object (big heavy = 0.08 ± 0.04) compared to observing lifts of the light one (small light = −0.09 ± 0.04; *p = 0.01)*. Interestingly, this analysis did not show significance for the main effect of GROUP as well as for its interaction with CUBE *(both F < 0.03, both p > 0.28, both η*^*2*^_*p*_ *< 0.04)*. As such, these findings further substantiate that in both the control and pSTS group, CSE modulation during lift observation was driven by the object’s actual weight (Figures 3 and 4).

Moreover, we explored whether CSE was still modulated by object weight after virtual lesioning of DLPFC using the same analysis as described in the preceding paragraph [GLM with between-factor GROUP (control and DLPFC) and within-factor CUBE (big heavy and small light)]. Briefly, this analysis failed to reveal significance for any of the main effects (GROUP and CUBE; *both F < 0.84, both p > 0.37, both η*^*2*^_*p*_ *< 0.03*) as well as their interaction effect *(F = 3.57, p = 0.06, all η*^*2*^_*p*_ *= 0.11)*. It is important to note that in the first paragraph of this results section (*‘corticospinal excitability during the experimental task’*), we already demonstrated for the control group that CSE modulation during lift observation was driven by object weight. Accordingly, considering that the interaction effect GROUP × CUBE was borderline significant and that the DLPFC group is included in this analysis, we decided to perform a final ANOVA_RM_ on the DLPFC group only with one within-factor CUBE (big heavy and small light). This was done to investigate whether CSE modulation in the DLPFC group was driven by CUBE. This analysis failed to show significance for CUBE *(F*_*(1,11)*_ *= 0.54, p = 0.48, η*^*2*^_*p*_ *= 0.05)*. In conclusion, these analyses provide no evidence at all that CSE was modulated during lift observation when DLPFC was virtually lesioned.

To end, we investigated whether CSE was modulated differently during lift observation and planning for the DLPFC and pSTS groups using a GLM with between-factor GROUP and within-factors ACTION (observation or planning), SIZE and WEIGHT. Interestingly, this analysis showed that CSE was significantly facilitated when observing or planning lifts of the heavy objects (heavy objects = 0.03 ± 0.02) compared to of the light ones (light objects = −0.05 ± 0.02; *p = 0.02) (main effect of WEIGHT: F*_*(1,22)*_ *= 6.68, p = 0.02, η*^*2*^_*p*_ *= 0.23)*. However, this WEIGHT effects was likely driven by the pSTS group as the significant interaction effect GROUP × WEIGHT *(F*_*(1,22)*_ *= 5.66, p = 0.03, η*^*2*^_*p*_ *= 0.20)* revealed that WEIGHT drove CSE modulation in the pSTS (heavy objects = 0.06 ± 0.02; light objects = −0.08 ± 0.03; *p = 0.01)* but not in the DLPFC group (heavy objects = −0.00 ± 0.02; light objects = −0.01 ± 0.03; *p = 1.00)*. In its turn, the significant difference between CSE modulation by the heavy and light objects for the pSTS group (GROUP × WEIGHT) was likely driven by the triple interaction effect GROUP × ACTION × WEIGHT *(F*_*(1,22)*_ *= 4.31, p = 0.05, η*^*2*^_*p*_ *= 0.16)*. Post-hoc exploration of this significant interaction effect revealed that, for the pSTS group, CSE was significantly facilitated during lift observation of the heavy objects (heavy objects = 0.11 ± −0.03) compared to of the light ones (light objects = −0.12 ± 0.03; *p = 0.04)* whereas this difference was absent during planning (heavy objects = 0.02 ± 0.04; light objects = −0.04 ± 0.04; *p = 1.00)*. In conclusion, these findings provide no evidence that CSE was modulated in the pSTS and DLPFC groups during lift planning (Figure 5). As we have no ‘control conditions’ (group without virtual lesioning during lift planning), these findings cannot be further interpreted.

**Figure 5.**
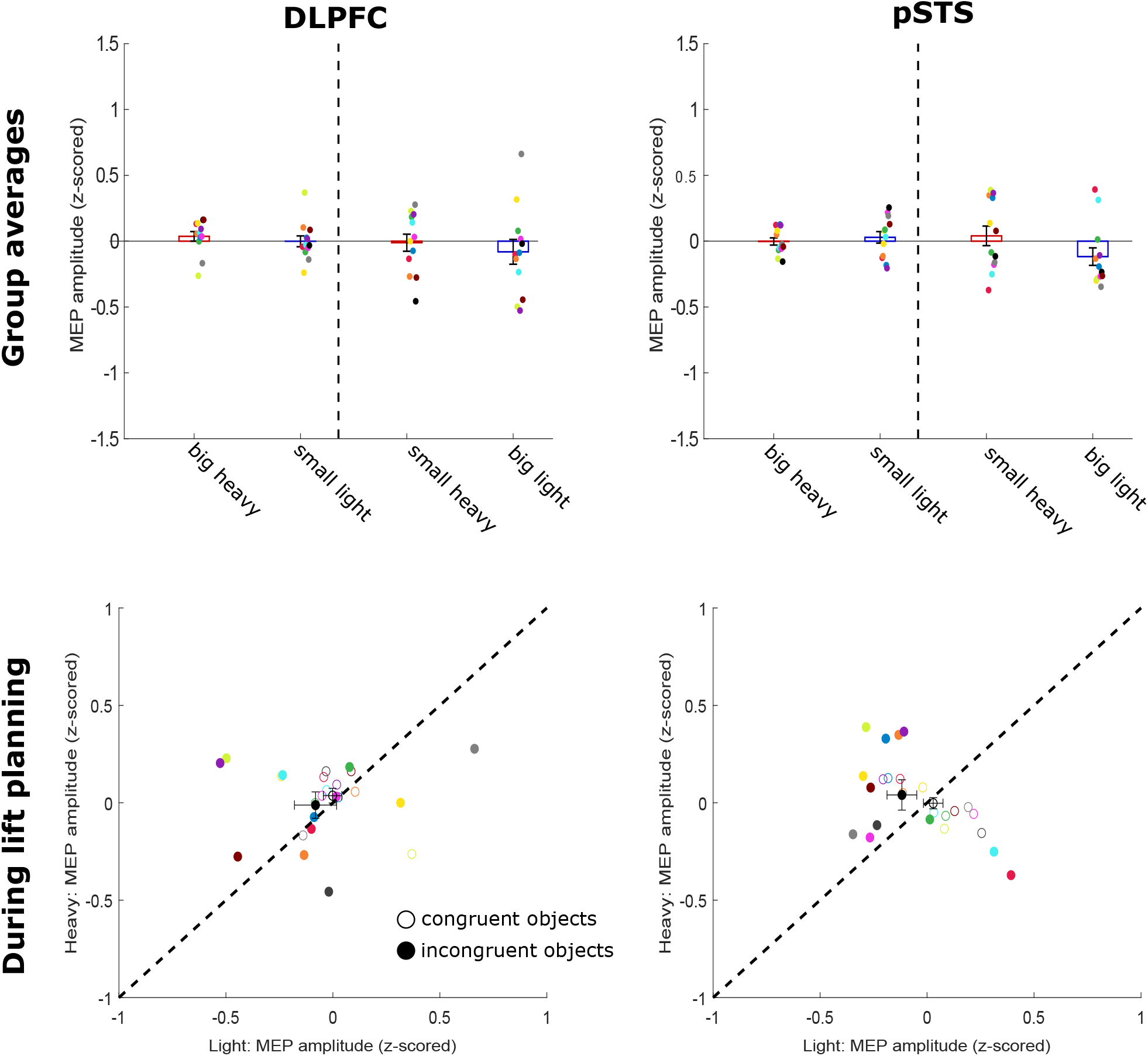
Modulation of corticospinal excitability during lift planning in the DLPFC and pSTS groups. **Top row.** Average MEP values (z-score) during lift planning pooled across participants for the DLPFC (left) and pSTS group (right). Left and right of the dashed line on each figure represent the congruent (big heavy and small light) and incongruent objects (small heavy and big light) respectively. Red and blue indicate heavy and light weights respectively. **Bottom row.** 2D visualization of the average MEP values (z-scored) during lift planning. MEP values for the heavy and light objects are shown on the y- and x-axis respectively. Each participant within each group is represented by two same colored circles (scatter). ‘Empty circles’ represent the congruent objects (MEP values for big-heavy on the y-axis and for small-light on the x-axis), ‘filled circles’ represent the incongruent objects (MEP values for small-heavy on the y-axis and for big-light on the x-axis). Black circles represent the group average ± SEM for the respective conditions. The black dashed line represents the equation y = x and indicates the line of ‘no CSE modulation’. Accordingly, scatter circles above or below the dashed line indicate that CSE, when observing lifts of planning heavier objects, was increased or decreased respectively. No intra-group significant differences are shown on the middle and bottom row.

To sum up, our results demonstrate that when participants only interact with objects having a congruent size-weight relationship (i.e. big-heavy or small-light), CSE during lift observation is modulated by the object weight as indicated by the size and/or the movement kinematics (control group). Interestingly, when objects with incongruent size-weight relationship (i.e. big light and small heavy) were included (baseline group), weight-driven modulation of CSE was ‘suppressed’ and CSE was modulated by ‘object congruency’ instead. That is, CSE was facilitated during observed lifting of objects with incongruent properties compared to of objects with congruent properties.

Moreover, our results also highlighted that virtual lesioning of pSTS abolishes the suppressive mechanism monitoring the observer’s weight expectations and restores weight-driven modulation of CSE during lift observation. As such, our results provide evidence for the causal involvement of pSTS in modulating CSE by monitoring the observer’s weight expectations during the observation of hand-object interactions. In addition, virtual lesioning of DLPFC eradicated both the suppressive mechanism as well as weight-driven motor resonance: During lift observation, we found no evidence that CSE was modulated at all. Accordingly, these findings suggest that DLPFC is causally involved in a ‘general’ modulation of CSE during the observation of hand-object interactions. To end, we did not find significant differences between the DLPFC and pSTS groups for lift planning. Considering that we have no ‘control’ group to compare with, these findings cannot be further interpreted.

### Force parameters of the participants

As mentioned before, we pooled all data with respect to factors related to TMS timing as preliminary analyses revealed that predictive force planning of the participants was not altered by single pulse TMS.

#### Normalized peak grip force rates

For both the group comparisons on the congruent objects only (all four groups) and on the objects with both congruency types (baseline, DLPFC and pSTS groups) neither the main effect of GROUP nor any of its interactions effects were significant (*all F < 0.86, all p > 0.47, all η*^*2*^_*p*_ *< 0.04)*.

First, for only the congruent objects these findings suggest that there is no evidence that the experimental groups scaled their grip forces (i.e. peak GFr values) differently, irrespective of whether they were exposed to only congruent object (control group) or to both congruent and incongruent objects (baseline, DLPFC and pSTS groups). Second, these findings also provide no evidence that virtual lesioning of either DLPFC or pSTS (DLPFC and pSTS groups) affected predictive grip force scaling based on lift observation compared to receiving no virtual lesioning (control and baseline groups). Aside from these results, all groups increased their grip forces significantly faster for the big heavy cube (big heavy = 0.48 ± 0.03) than for the small light one (small light = −0.43 ± 0.03) (*main effect of CUBE: (F*_*(1,55)*_ *= 353.70, p < 0.001, η*^*2*^_*p*_ *= 0.87)*. All group averages are shown in Figure 6.

**Figure 6.**
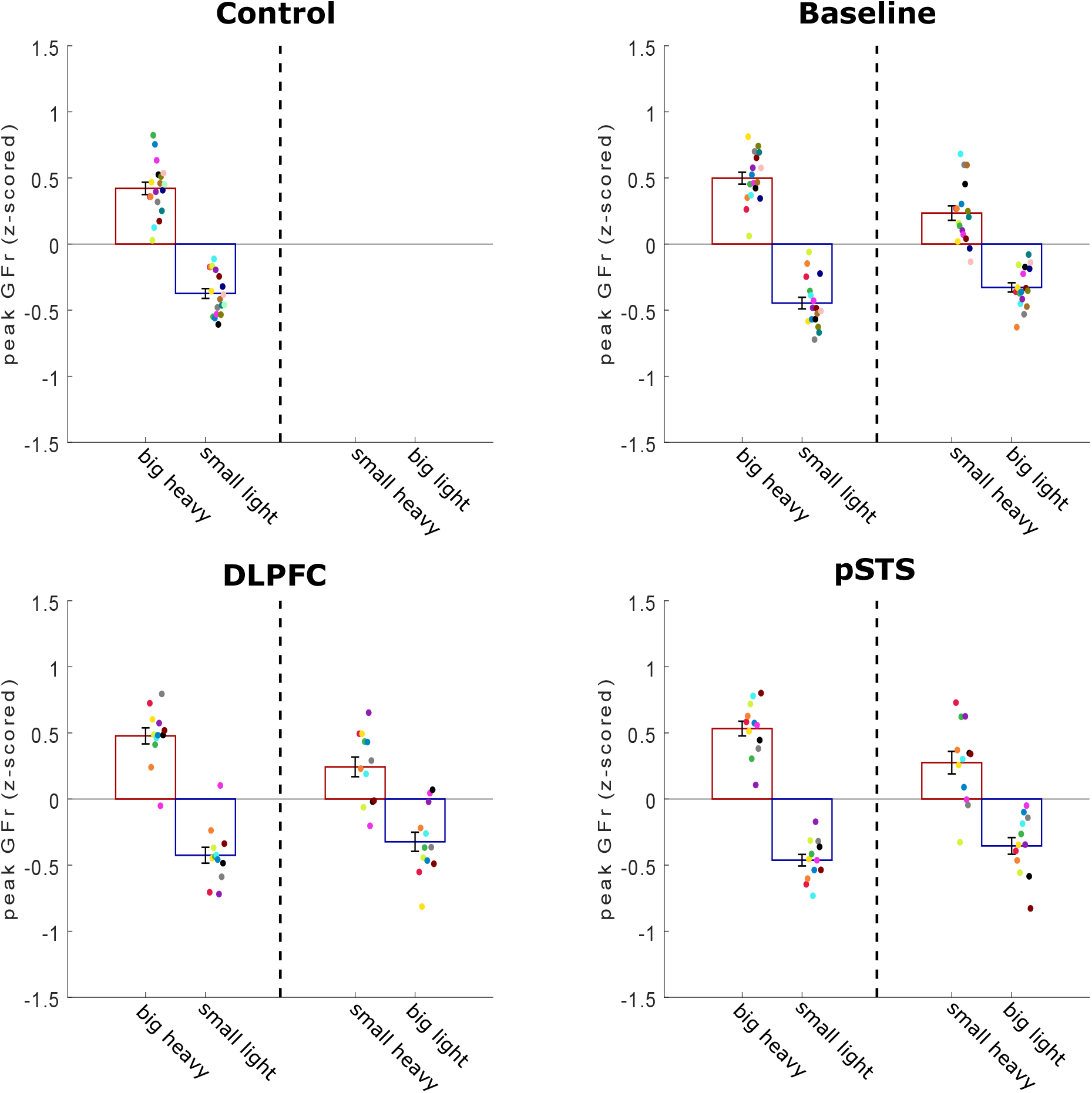
Peak grip force rates of the participants. Average peak grip force rate (GFr) value (z-scored) for each group separately. Left and right of the dashed line on each figure represent the congruent (big heavy and small light) and incongruent objects (small heavy and big light), respectively. Within each experimental group, each colored circle (scatter) represents the average peak GFr value for one participant in that specific condition. All data is presented as the mean ± SEM. No intra-group significant differences are shown on this figure.

Moreover, these findings are similar for the groups that interacted with both congruent and incongruent objects. That is, the baseline, DLPFC and pSTS groups increased their grip forces significantly faster for the heavy objects (heavy = 0.38 ± 0.03) than for the light ones (light = −0.39 ± 0.02; *p < 0.001*) *(main effect of WEIGHT: (F*_*(1,38)*_ *= 255.93, p < 0.001, η*^*2*^_*p*_ *= 0.87)*. However, although these groups were able to scale their grip forces to the actual object weight, they were still biased by the size as they increased their grip forces significantly faster for the big objects (big objects = 0.08 ± 0.02) than for the smaller ones (small objects = −0.10 ± 0.02; *p < 0.001) (main effect of SIZE: (F*_*(1,38)*_ *= 23.69, p < 0.001, η*^*2*^_*p*_ *= 0.38)*. Lastly, post-hoc analysis of the significant interaction effect WEIGHT × SIZE *(F*_*(1,38)*_ *= 5.42, p = 0.025, η*^*2*^_*p*_ *= 0.12)* highlighted that these groups also increased their grip forces significantly faster for the big heavy object (big heavy = 0.50 ± 0.03) than for the small heavy one (small light = 0.25 ± 0.04; *p < 0.001)*. This difference was absent for the light objects (small light = −0.44 ± 0.03; big light = −0.34 ± 0.03; *p* = 0.08).

#### Normalized peak load force rates

The findings for peak LFr were nearly identical to those for peak GFr. Indeed, for both comparisons [congruent objects only: all groups; both congruent and incongruent objects: baseline, DLPFC and pSTS groups], the main effect of GROUP as well as all its interactions effects were not significant (*all F < 0.72, all p > 0.49, all η*^*2*^_*p*_ *< 0.04)*. Accordingly, we did not find any evidence that predictive load force planning based on lift observation was affected by (1) the presence of the incongruent objects (control group vs baseline, DLPFC and pSTS groups) (2) or by the virtual lesioning of DLPFC or pSTS (control and baseline groups vs DLPFC and pSTS groups). Similar to our findings for peak GFr, participants increased their load forces significantly faster for the big heavy cube (big heavy = 0.42 ± 0.02) than for the small light one (small light = −0.39 ± 0.02; *p < 0.001)* (*main effect of CUBE: (F*_*(1,55)*_ *= 339.57, p < 0.001, η*^*2*^_*p*_ *= 0.86)*.

Again, the baseline, DLPFC and pSTS groups, that interacted with both congruent and incongruent objects, increased their load forces significantly faster for the heavy objects (heavy = 0.35 ± 0.02) than for the light ones (light = −0.35 ± 0.2; *p < 0.001)* (*main effect of WEIGHT: (F*_*(1,38)*_ *= 304.80, p < 0.001, η*^*2*^_*p*_ *= 0.89)* although they were also biased by object size (big: peak LFr = 0.05 ± 0.02; small: peak LFr = −0.05 ± 0.02; *p = 0.004)* (*main effect of SIZE: (F*_*(1,38)*_ *= 9.10, p = 0.005, η*^*2*^_*p*_ *= 0.19)*. All group averages are shown in Figure 7 without intra-group significant differences being shown.

**Figure 7.**
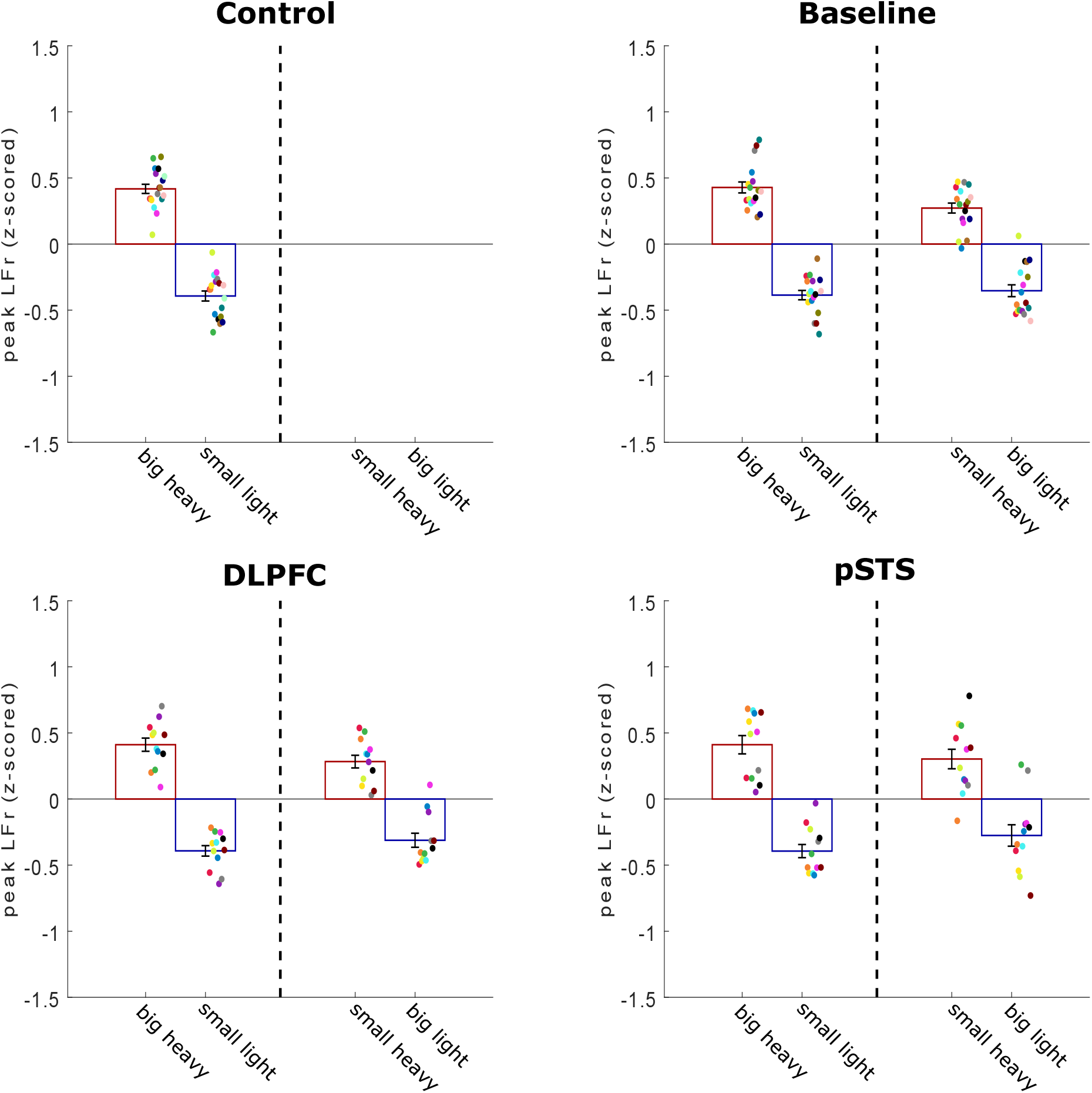
Peak load force rates of the participants. Average peak load force rate (LFr) values (z-scored) for each group separately. Left and right of the dashed line on each figure represent the congruent (big heavy and small light) and incongruent objects (small heavy and big light), respectively. Within each experimental group, each colored circle (scatter) represents the average peak LFr value for one participant in that specific condition. All data is presented as the mean ± SEM. No intra-group significant differences are shown on this figure.

#### Normalized loading phase duration

Our findings for the participants’ loading phase duration were identical to those for peak GFr: For congruent objects only (all groups) and the congruent and incongruent objects combined (baseline, DLPFC and pSTS groups) our analyses did not show significance for the main effect of GROUP as well as its interaction effects (*all F < 2.07, all p > 0.140, all η*^*2*^_*p*_ *< 0.10)*, again suggesting that our experimental groups did not differ significantly from each other. Again, the GLM for the congruent objects only showed that the main effect of CUBE was significant *(F*_*(1,55)*_ *= 2717.64, p < 0.001, η*^*2*^_*p*_ *= 0.90)* indicating that all groups lifted the big heavy object (big heavy = 0.83 ± 0.02) slower than the small light one (small light = −0.80 ± 0.02; *< 0.001*).

In line with our peak GFr findings, the groups (baseline, DLPFC and pSTS), interacting with both congruent and incongruent objects lifted the heavy objects (heavy = 0.91 ± 0.03) significantly slower than the light ones (light = −0.80 ± 0.02; *p < 0.001*) *(main effect of WEIGHT: F*_*(1,38)*_ *= 1139.85, p < 0.001, η*^*2*^_*p*_ *= 0.97)* although they were still biased by the object size as they lifted the big objects faster than the small ones (big = 0.01 ± 0.01; small = 0.09 ± 0.02; *p < 0.001) (main effect of SIZE: F*_*(1,38)*_ *= 18.43, p < 0.001, η*^*2*^_*p*_ *= 0.33)*. Finally, post-hoc analysis of the significant interaction effect WEIGHT × SIZE (*F*_*(1,38)*_ *= 23.33, p < 0.001, η*^*2*^_*p*_ *= 0.38)* revealed that all groups lifted the big heavy object (big heavy = 0.82 ± 0.02) significantly faster than the small heavy one (small heavy = 0.99 ± 0.04; *p < 0.001*) although this difference was absent for the light objects (small light = −0.81 ± 0.02; big light = −0.80 ± 0.03; *p = 1.00)*. All group averages are shown in Figure 8 without intra-group significant differences being shown.

**Figure 8.**
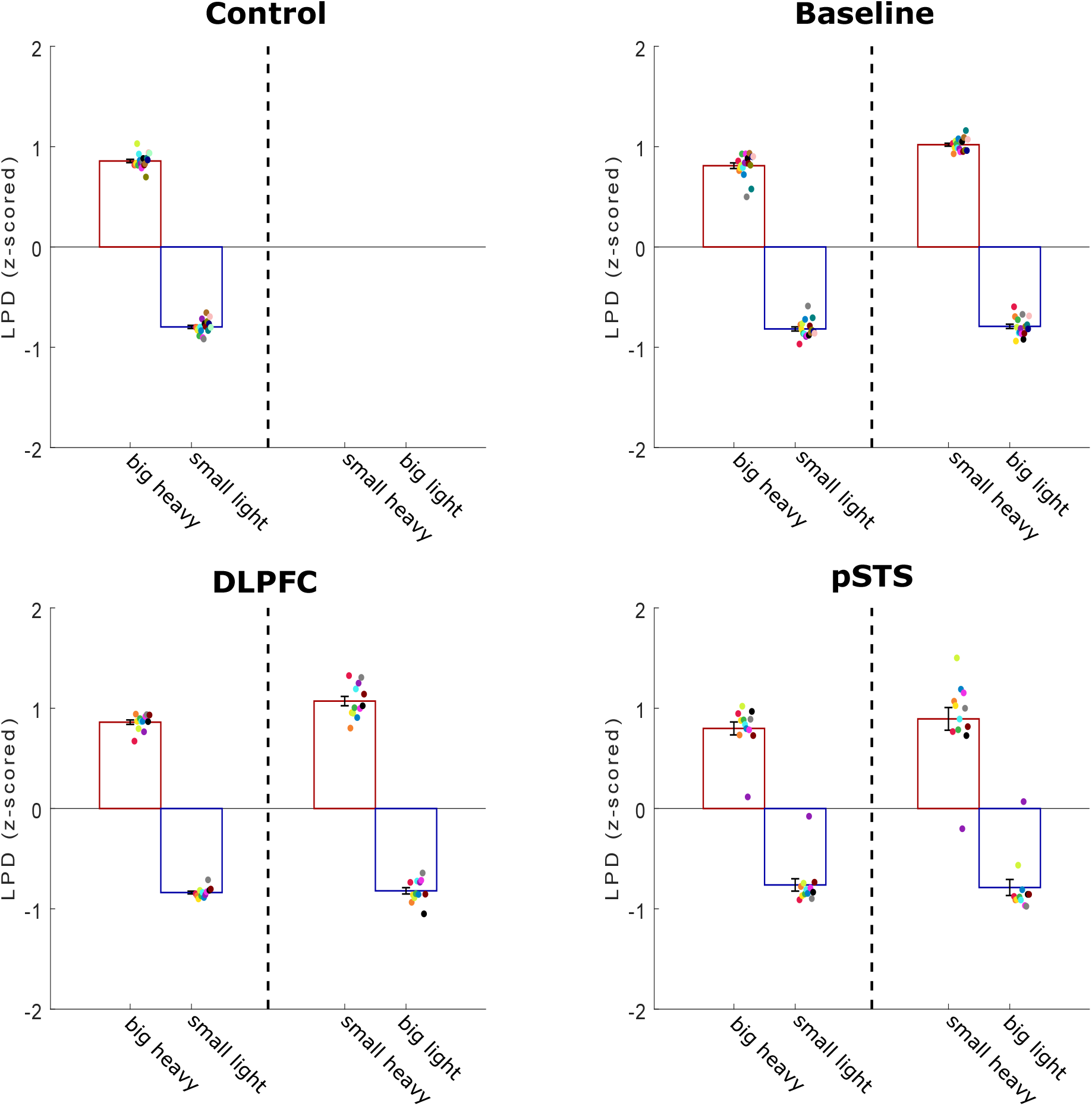
Loading phase duration of the participants. Average loading phase duration (LPD) values (z-scored) for each group separately. Left and right of the dashed line on each figure represent the congruent (big heavy and small light) and incongruent objects (small heavy and big light), respectively. Within each experimental group, each colored circle (scatter) represents the average peak LPD value for one participant in that specific condition. All data is presented as the mean ± SEM. No intra-group significant differences are shown on this figure.

To sum up, participants lifted the objects [SIZE: big or small by WEIGHT: heavy or light] in turns with the actor and were instructed that the object in their trial was always identical, both in terms of size and weight, to the object the actor lifted in the previous trial. As such, participants could potentially rely on lift observation to estimate object weight and plan their own lifts accordingly. Importantly, our results support this notion: In line with Rens and Davare (2019), our results demonstrate that the groups who interacted with both the congruent and incongruent objects were able to detect the incongruent objects based on observed lifts performed by the actor. Indeed, our findings for the baseline, DLPFC and pSTS groups showed that subjects scaled their fingertip forces to the actual weight of the incongruent objects (main effect of WEIGHT). However, it is important to note that these groups were still biased by object size as, on average, subjects scaled their fingertip forces faster for the large objects than for the small ones (main effect of SIZE). Moreover, exploration of the significant interaction effect of WEIGHT × SIZE for peak GFr and LPD indicated that this effect was primarily driven by the significant difference between heavy objects. Lastly, considering that we did not find significant differences between the baseline group on one side and the DLPFC and pSTS groups on the other side shows that virtual lesioning of either region did not affect predictive lift planning based on lift observation. As such, our findings related to the force parameters indicate that DLPFC and pSTS are not causally involved in neither weight perception during lift observation nor in updating the motor command based on lift observation.

### Force parameters of the actor

#### Normalized peak grip force rates

Comparing the congruent objects only across all four groups, the actor increased his grip forces significantly faster for the big heavy object (big heavy = 0.8 ± 0.02) than for the small light one (small light = −0.79 ± 0.01; *p < 0.001*) *(main effect of WEIGHT*: *F*_*(1,55)*_ *= 3328, p < 0.001, η*^*2*^_*p*_ *= 0.98).* Although the main effect of group was not significant, the interaction effect of GROUP × CUBE *(F*_*(3,55)*_ *= 5.85, p = 0.002, η*^*2*^_*p*_ *= 0.24)* was. Post-hoc analysis of this interaction effect showed that the actor scaled his grip forces significantly faster for the big heavy object in the baseline group (baseline: big heavy = 0.89 ± 0.03) compared to the control group (control: big heavy = 0.76 ± 0.03, *p = 0.02*). However, all other between-group differences in the actor’s lifting performance for the big heavy object were not significant (DLPFC: big heavy = 0.88 ± 0.04; pSTS: big heavy = 0.78 ± 0.03; *all p > 0.12)*. Conversely, this was identical for the small light object with the actor scaling his grip forces significantly slower for the small light object in the baseline group (baseline: small heavy = −0.84 ± 0.02) than in the control group (control: small heavy = −0.72 ± 0.02; *p = 0.05)*. Again, all other between-group actor differences for the small light object were not significant (DLPFC: small light = −0.83 ± 0.03; pSTS: small light = −0.76 ± 0.03; *all p > 0.24)*.

For the comparisons including the incongruent objects (baseline, DLPFC and pSTS groups), it is important to note that the interaction effect SIZE × WEIGHT (*F*_*(1,38)*_ *= 5.52, p = 0.02, η*^*2*^_*p*_ *= 0.13)* was significant. Post-hoc analysis showed that the actor increased his grip forces similarly for the light objects (small light = −0.81 ± 0.02; big light = −0.83 ± 0.03; *p = 1.00)* but not for the heavy ones (big heavy = 0.85 ± 0.02; small heavy = 0.79 ± 0.04; *p = 0.03)*. As our results indicate that the actor increased his grip forces slower for the small heavy object compared to the big heavy object suggesting that he was biased by the object’s size during his own trials.

#### Normalized peak load force rates

In line with our findings for grip force rates, the actor increased his load forces significantly faster for the big heavy cuboid (big heavy = 0.80 ± 0.02) than the small light one (small light = −0.72 ± 0.02; *p < 0.001*) (congruent objects only: *main effect of CUBE: F*_*(1,,55)*_ *= 1950.87, p < 0.001, η*^*2*^_*p*_ *= 0.97)*. Importantly, post-hoc exploration of the significant interaction effect GROUP × CUBE *(F*_*(3,55)*_ *= 3.87, p = 0.01, η*^*2*^_*p*_ *= 0.17)*, did not reveal any relevant significant differences in the actor’s performance between groups on the big heavy object (control = 0.71 ± 0.04; baseline = 0.84 ± 0.04; DLPFC = 0.85 ± 0.04; pSTS = 0.79 ± 0.04; *all p > 0.18)* or the small light one (control = −0.63 ± 0.03; baseline = −0.76 ± 0.03; DLPFC = −0.76 ± 0.04; pSTS = −0.71 ± 0.04; *all p > 0.18)*.

However, the analysis on both the congruent and incongruent objects, showed that the actor scaled his load forces differently based on object size for both the light objects (small light = −0.74 ± 0.02; big light = −0.82 ± 0.03; *p = 0.05)* and the heavy ones (big heavy = 0.83 ± 0.03; small heavy = 0.74 ± 0.04; *p = 0.04) (SIZE × WEIGHT: F*_*(1,,38)*_ *= 15.40, p < 0.001, η*^*2*^_*p*_ *= 0.29)*. Finally, it is important to note that neither the main effect of GROUP nor its interaction effects were significant *(all F < 1.03, all p > 0.37, all η*^*2*^_*p*_ *< 0.5).* As such, we did not find evidence that the actor scaled his load forces differently for the different experimental groups.

#### Normalized loading phase duration

Comparing only the congruent objects across all four groups showed that LPD of the actor was significantly longer when lifting the big heavy object (big heavy = 0.76 ± 0.02) than the small light one (small light = −0.85 ± 0.02; *p < 0.001)* (congruent objects only: *main effect of CUBE: F*_*(1,55)*_ *= 2883.95, p < 0.001, η*^*2*^_*p*_ *= 0.98)*. For the comparison on both the congruent and incongruent objects, the interaction effect SIZE × WEIGHT *F*_*(1,38)*_ *= 57.40, p < 0.001, η*^*2*^_*p*_ *= 0.60)* was significant. Critically, the post-hoc analysis revealed that the actor lifted the small objects significantly slower than the big ones. That is, the LPD when lifting the big heavy object (big heavy = 0.76 ± 0.02) was significantly shorter than when lifting the small heavy one (small heavy = 0.89 ± 0.03; *p < 0.001)*. Accordingly, this significant difference was also present for the light objects (small light = −0.84 ± 0.02; big light = −0.68 ± 0.02; *p < 0.001)*. Although these findings suggest that the actor’s lifting speed was biased by object size, he still lifted the light objects significantly faster than the heavy ones *(SIZE × WEIGHT: all p < 0.001)*.

In sum, these findings indicate that, in general, the actor scaled his fingertip forces towards the actual object weight for both the congruent and incongruent objects. However, it is important to note that the actor was biased by object size when interacting with the incongruent objects. Across all groups (except the control group which did not interact with the incongruent objects), the actor increased his fingertip forces faster for the big than for the small objects, resulting in a shorter LPD for the larger objects. Presumably, as participants were able to lift the objects (of which they could only predict object weight by relying on the actor’s lifting) skilfully, it is plausible that these found differences in the actor’s lifting performance drove the participants’ ability to estimate object weight during observed lifting. Accordingly, these differences in observed lifting performance should also have driven modulation of CSE. Finally, except for one difference for normalized grip force rates, the actor scaled his fingertip forces similarly across all groups. Importantly, these findings substantiate that our inter-group differences, with respect to CSE modulation, are not driven by differences in the actor’s lifting performance between groups but rather by experimental set-up differences [presence of incongruent objects vs. only congruent objects; virtual lesioning of pSTS or DLPFC vs. no virtual lesion].

## Discussion

First, we investigated how CSE is modulated during observation of lifting actions (i.e. ‘motor resonance’). Our control experiment findings align with previous literature (Alaerts et al., 2010a, 2010b): When participants observed lifts of objects with a congruent only size-weight relationship, CSE was modulated by object weight. However, our baseline group findings highlight that weight-driven motor resonance effects are easily suppressed when weight cannot be reliably predicted based on size: When participants observed lifts of objects with congruent and incongruent size-weight relationships, CSE was larger when observing lifts of incongruent objects, regardless of their size and weight. Interestingly, this suggests that ‘typical’ weight-driven motor resonance was suppressed by a mechanism monitoring size-weight congruence. However, we found these differences at different time points during action observation (Figure 3), indicating that the baseline group perceived the small-light object weight *before lift-off*. Presumably, participants estimated weight based on the actor’s reaching phase as Eastough and Edwards (2007) demonstrated that an individual’s reaching phase depends on the object’s mass. However, we cannot substantiate this assumption as we did not record the actor’s reaching phase. Finally, in line with Rens and Davare (2019), the baseline group was able to generate the appropriate fingertip force scaling to lift the objects skillfully after lift observation.

Second, we investigated the causal involvement of top-down inputs in the suppressive mechanism monitoring size-weight congruence by disrupting either pSTS or DLPFC using cTBS. Strikingly, pSTS virtual lesions abolished the suppressive mechanism and restored weight-driven motor resonance suggesting that pSTS is pivotal in monitoring weight expectations during lift observation. In contrast, DLPFC virtual lesions eradicated all modulation of motor resonance suggesting that DLPFC is causally involved in the overall modulation of motor resonance. Although virtual lesions of DLPFC and pSTS altered motor resonance, we found no evidence that predictive lift planning, after lift observation, was affected. This suggests that adequate motor planning is not necessarily related to motor resonance effects.

Regarding our baseline group, Alaerts et al. (2010b) showed that, when participants observed lifts of objects with incongruent properties, motor resonance was still driven by weight as cued by the movement kinematics. Our results contrast theirs by showing that motor resonance was rather driven by size-weight congruence. Critically, our study differs from theirs on three major points. First, participants in their study did not manipulate the objects. Second, their participants were not required to respond after observation (verbally or behaviorally) and third, whereas we used a skewed proportion of congruent and incongruent trials, they used equal proportions.

It is unlikely that our baseline group findings are entirely driven by the skewed proportion of congruent vs. incongruent trials: Pezzetta et al. (2018) demonstrated using electroencephalography that participants elicit typical error-monitoring activity whether larger or smaller proportions of erroneous grasping are observed. A plausible explanation, that our findings are driven by the experimental context rather than the skewed proportion, resides in another study of Alaerts et al. (2012): They demonstrated that motor resonance can reflect object weight predictively *during observed reaching* and, thus, when the actual object weight cannot yet be veridically identified. However, they used a blocked design and never challenged the participants’ expectations. Thus, in our baseline group, randomly inserting trials with incongruent object size-weight properties might have caused a top-down mechanism to suppress weight-driven motor resonance. Arguably, this mechanism might be useful to prevent motor resonance from encoding object weight based on an incorrect prediction. That is, when a mismatch between expected and actual object weight is identified, this top-down mechanism releases all suppression allowing a sudden increase in CSE, which signals that the motor command will need to be updated from the one initially predicted based on object size, to the correct one based on the actor’s lifting kinematics. As such, the contextual importance of accurately estimating object weight during observation might have driven this mechanism to suppress weight-driven motor resonance.

Motor resonance has been argued to rely on the putative human mirror neuron system (hMNS). First discovered in monkeys (di Pellegrino et al. 1992), mirror neurons are similarly activated when executing or observing the same action and have been argued to be involved in action understanding by ‘mapping’ observed actions onto the cortical representations involved in their execution (Cattaneo and Rizzolatti 2009). The hMNS is primarily located in M1, ventral premotor cortex (PMv) and anterior intraparietal area (AIP) (Rizzolatti et al. 2014). Importantly, these regions also constitute the cortical grasping network which is pivotal in planning and executing grasping actions (for a review see: Davare et al., 2011) further substantiating hMNS’ involvement in action understanding.

However, Amoruso and Finisguerra (2019) argued that motor resonance only reflects an automatic replica of observed actions, if observed in isolation, but that it can be modulated by top-down inputs in presence of contextual cues. Our results support this hypothesis: Weight-driven motor resonance was present when weight expectations were never challenged (control group), but turned out to be suppressed when a size-weight mismatch was introduced (baseline group). Although we demonstrated a systematic effect of size-weight contingency on motor resonance, Figure 3 shows that the presence of incongruent trials (baseline group, right) also led to a larger between-subject variability compared to the control group (Figure 3, left). This might be explained by the baseline group subjects relying on different strategies to extract weight-related information: either focusing on the movement kinematics or the size-weight contingency (Amoruso and Finisguerra 2019).

In our second experiment, we investigated the origins of the suppressive mechanism and found that disrupting pSTS restores weight-driven motor resonance, suggesting that pSTS is causally involved in monitoring expectations during observation. These findings are plausible as pSTS is crucial in perceiving biological motion (Grossman et al., 2005), which is indicative of object weight (Hamilton et al. 2007), and in monitoring execution errors during observation (Pelphrey et al., 2004). Although pSTS does not contain mirror neurons (Hickok, 2009, 2013) and shares no connections with M1 (Iacoboni, 2005; Nelissen et al., 2011), it accesses the putative hMNS through reciprocal connections with AIP (Galletti and Fattori, 2017; Nelissen et al., 2011). Plausibly, pSTS modulates CSE through AIP-PMV and PMv-M1 connections (Davare et al., 2011; Gerbella et al., 2017). Indeed, our results suggest that pSTS monitors weight expectations during observed lifting and masks typical motor resonance effects when expectations can be incorrect. Plausibly, virtual lesioning of pSTS abolishes expectation-related input to AIP, restoring the automatic mapping of observed movement features. In addition, when expectations are never tested (control group), pSTS might not provide this top-down input and does not mask weight-driven motor resonance. However, future research is necessary to substantiate the latter.

We also investigated the causal involvement of DLPFC in monitoring weight expectations: Our results show that disrupting DLPFC eradicated both the expectation monitoring mechanism and also weight-driven motor resonance arguing that DLPFC is pivotal in the overall modulation of CSE during lift observation, irrespective of the underlying mechanism. Our results align with those of Ubaldi et al. (2015): They showed that when motor resonance effects were altered by a visuomotor training task, the trained resonance could be eradicated by virtual lesioning of DLPFC, suggesting that DLPFC is critical in modulating rule-based motor resonance. Importantly, our results extend on theirs by demonstrating that virtual lesioning of DLPFC eradicates not only trained effects but also effects which are considered to be automatic. It is plausible that DLPFC can modulate motor resonance: Although DLPFC does not contain mirror neurons (Hickok 2009, 2013), it is reciprocally connected with PMv (Badre and D’Esposito 2009) and involved in action observation and processing contextual information (Raos and Savaki 2017; Rozzi and Fogassi 2017).

A limitation of the present study is that we used one TMS timing in the virtual lesion groups, due to time constrains. We only probed motor resonance after observed lift-off as we found the strongest effects of the suppressive mechanisms for our baseline group at this timing. In addition, Ubaldi et al. (2015) demonstrated that motor resonance driven by visuomotor associations is only altered during late but not early movement observation. Therefore, it seemed valid to focus on this timing. A second limitation concerns the absence of sham cTBS in experiment 2. Noteworthy, virtual lesioning of DLPFC and pSTS modulated CSE differently, indicating that the stimulation site was relevant. However, probing motor resonance when observing lifts of congruent objects only, combined with cTBS disruption of DLPFC and pSTS, could further substantiate our findings. The last limitation is that we did not use a within-subject design. Considering our hypothesis that individuals’ expectations alter motor resonance, we opted for a between-subject design to ensure all participants have the same expectations when performing the behavioral task (for the first time).

In conclusion, the present study shows that motor resonance is not robust but influenced by contextual differences. We argue that motor resonance should be carefully interpreted in light of the hMNS putative roles. Our results indicate that bottom-up motor resonance effects, driven by observed movement features, can only be probed when top-down suppressive expectation monitoring mechanisms from pSTS are not triggered. Moreover, DLPFC is pivotal in the global modulation of CSE during action observation. Altogether, these findings shed new light on the theoretical framework in which motor resonance effects occur and overlap with other cortical processing essential for the sensorimotor control of movements.

## Acknowledgements

GR is a doctoral student funded by a Research Foundation Flanders (FWO) Odysseus Project (Fonds Wetenschappelijk Onderzoek, Belgium; grant: G/0C51/13N) awarded to MD. VVP is funded by an FWO post-doctoral fellowship (grant: 12X7118N). MAG was supported by FWO Research Foundation (Grants G099516N).

